# Signaling dynamics control cell fate in the early Drosophila embryo

**DOI:** 10.1101/342998

**Authors:** Heath E. Johnson, Stanislav Y. Shvartsman, Jared E. Toettcher

## Abstract

The Erk mitogen-activated protein kinase plays diverse roles in animal development, where its activity is associated with phenomena including cell migration, proliferation and differentiation. Its widespread reuse raises a conundrum: when a single kinase like Erk is activated, how does a developing cell know which fate to adopt? Here, we combine precise optogenetic control with genetic perturbations to dissect Erk-dependent cellular responses in the early Drosophila embryo. We find that light-stimulated Erk activity is sufficient to ‘posterior-ize’ the majority of the embryo, leading to massive apical constriction through expression of the autocrine receptor-ligand pair *mist* and *fog*. Ectopic contraction at non-terminal positions requires at least 1 h of high-amplitude Erk signaling, whereas a 30 min pulse of Erk activity patterns non-contractile neurogenic ectoderm during the same time window. In contrast to the canonical ‘transient versus sustained’ model, the cell fate switch is triggered by the cumulative load of Erk signaling, not the duration of a single persistent pulse. Our results reveal that the early fly embryo harbors a classic example of dynamic cell fate control, where the total dose of Erk activity selects between two distinct physiological outcomes.

---

One of the great mysteries of animal development is how a small number of intracellular signals can be reused at different positions and times to coordinate a wide range of cell fate decisions. A classic paradigm for this one-to-many mapping is the idea of a morphogen, a substance whose concentration varies with embryonic position and where different concentrations are sufficient to induce different cell fates^1^. Alternatively, cell fates may be specified by combinatorial control: the spatial overlap between particular combinations of patterning cues^2^. A third model, dynamic control, holds that a single signal could select among cellular responses based on features such as the amplitude, duration, or frequency of pathway activation^3^. Although all three paradigms have been proposed to explain cell fate decisions, directly demonstrating which underlies particular developmental decisions has been extremely challenging. Researchers typically lack the ability to vary a single feature, such as the concentration, duration, or spatial range of a signal, while holding others constant.

Here, we set out to dissect cell fate control in a model developmental context: the Erk-dependent control of cellular responses in the early *Drosophila* embryo. Two factors make the early embryo ideal for such a study. First, Erk activity is required for cells to adopt distinct fates at three different positions: the anterior pole, which forms head structures; the ventral midline, a stripe on each side of the embryo that forms a population of neuronal precursors; and the posterior pole, which forms endoderm and undergoes profound movements during gastrulation (Figure 1A). Second, Erk signaling can be precisely controlled in space and time using optogenetics, enabling the experimentalist to directly test how different signal features map to gene expression and cell fate responses^4^^-^^6^.

**Figure 1.**
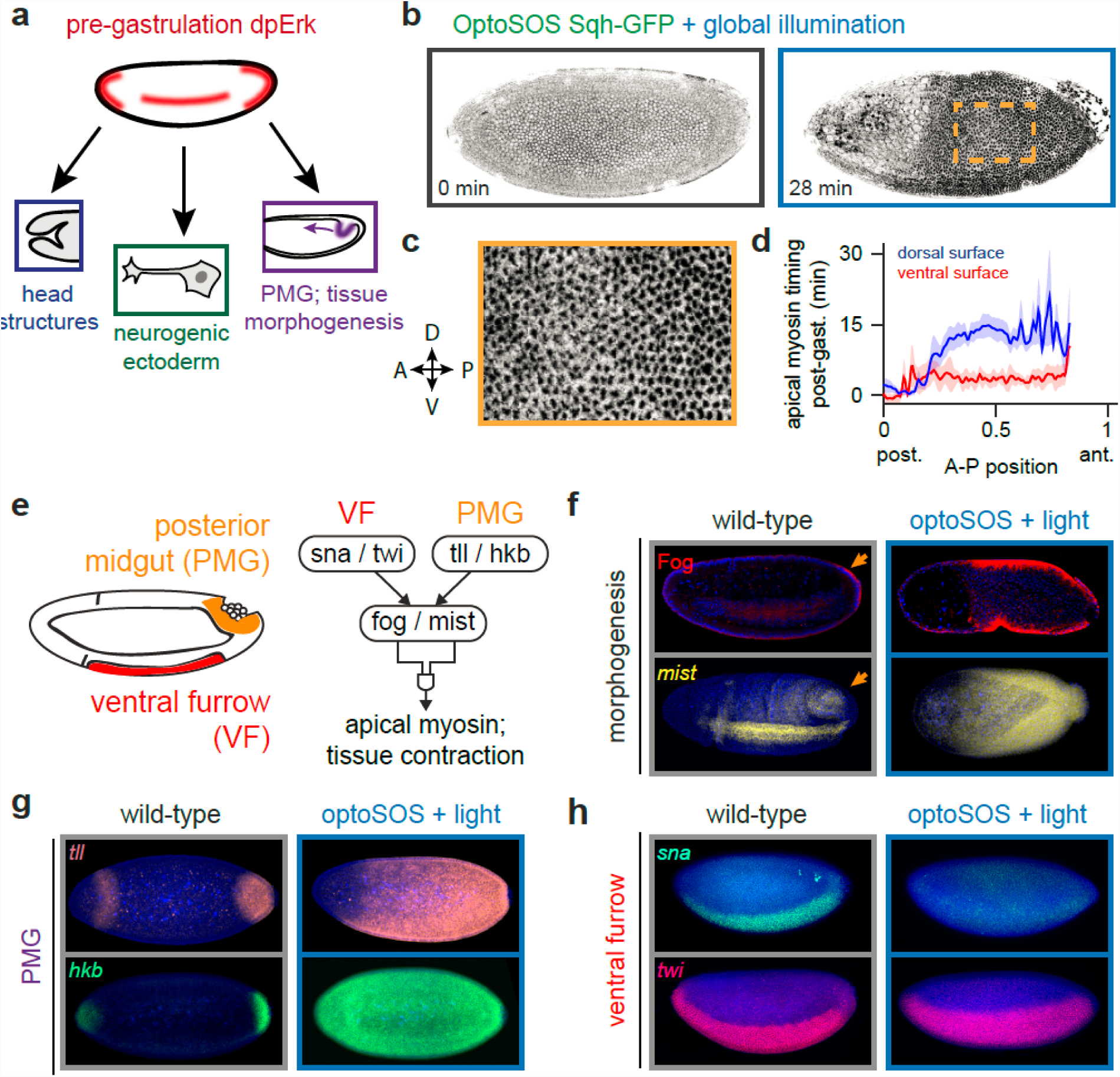
Erk signaling is sufficient for programming contractile fates in the early Drosophila embryo. (**a**) Erk activity is present at three locations within the blastula and coordinates distinct fates. Of these, only the posterior undergoes apical constriction and invagination during gastrulation. (**b**) Max-projected Sqh-GFP images from an OptoSOS embryo under continuous global illumination during NC14 (left) and after the start of gastrulation (right). Dark regions indicate apical myosin localization (See also Movie S2). (**c**) Detail of boxed region from **b**. (**d**) Time of initial apical myosin appearance as a function of A-P and D-V position from 5 individual OptoSOS embryos. (**e**) Schematic of the genetic network controlling tissue contractility in the ventral furrow and posterior midgut. (**f**) RNA FISH for *mist* and Fog immunostaining in gastrulating wild-type and OptoSOS embryos illuminated for 2 h prior to fixation. (**g-h**) RNA FISH for (**g**) *tll* / *hkb* and (**h**) *sna* / *twi* in wild-type and OptoSOS embryos illuminated for 2 h prior to fixation.

We previously showed that in OptoSOS-expressing fly embryos, stimulation with blue light activates Erk at least as strongly as the maximum level reached in wild-type embryos^6^, unlike prior gain-of-function mutants which only modestly increase embryo-wide Erk activity (Figure S1AC)^7^. Brightfield imaging of gastrulating embryos further revealed that Erk activation led to profound defects in morphogenesis, expanding the population of contractile cells and leading to large-scale tissue rearrangements^6^. Yet the precise nature of these defects has not yet been determined, leaving many questions open. Which populations of cells are susceptible to Erkinduced reprogramming into contractile tissue? What Erk-dependent genetic network drives contractility? Finally, if Erk is sufficient to induce morphogenesis, why is it that the anterior pole and ventral midline do not normally contract despite their high levels of Erk signaling?

To identify which cells could be reprogrammed by Erk to drive tissue morphogenesis, we first set out to image the dynamics of contraction at single-cell resolution in live OptoSOS embryos. We used a fluorescent myosin light chain, Sqh-GFP, which redistributes from the basal to apicala surface in invaginating cells during gastrulation^8, 9^. Dark-incubated OptoSOS/Sqh-GFP embryos gastrulated normally, exhibiting apical myosin redistribution in two tissues that normally invaginate, the ventral furrow and at the posterior pole (Movie S1; Figure S2A). In contrast, blue light stimulation of OptoSOS/Sqh-GFP embryos led to a massive expansion of apical myosin in the illuminated region (Figures 1B and S2B-D; Movie S2-3). We found that contractility was induced uniformly, with all cells exhibiting apical myosin localization within the contractile domain (Figure 1C). However, the timing of contraction varied with embryonic position, propagating in a posterior-to-anterior wave over 15 min, with positions on the ventral surface constricting approximately 10 min before their dorsal counterparts (Figure 1D). Interestingly, no apical myosin or contractility was observed in the anterior-most positions of the embryo, even under global illumination. Our data thus suggests that Erk is sufficient to trigger tissue contractility in the posterior 85% of the embryo, but that the timing of contraction is regulated along both the anterior-posterior (A-P) and dorsal-ventral (D-V) axes and set independently of Erk.

Which molecular pathways are activated by Erk to drive tissue contractility? We reasoned that Erk-induced contractility might reflect the ectopic formation of a tissue that normally contracts during gastrulation, the ventral furrow (VF) or posterior midgut endoderm (PMG) (Figure 1E). In both cases, tissue contractility is thought to be driven by the localized expression of *folded gastrulation* (*fog*) and *mist*, a secreted ligand and its cognate G protein coupled receptor (GPCR)^10, 11^. Although *fog* and *mist* are associated with both VF and PMG contraction, their expression is controlled by distinct upstream pathways; expression in the VF is regulated by *snail* (*sna*) and *twist* (*twi*) and in the PMG by *tailless* (*tll*) and *huckebein* (*hkb*).

Staining for Fog protein and *mist* mRNA in gastrulating, light-stimulated OptoSOS embryos revealed that their expression exactly overlapped the light-stimulated contractile domain, with much lower expression at far anterior positions (Figure 1F). Staining for upstream pathway markers revealed that *fog*/*mist* expression is likely driven by PMG and not VF gene networks: *tll* expression expanded over the contractile domain, with *hkb* expression extending even further to the anterior pole (Figure 1G). In contrast, *twi* expression was unaffected and *sna* expression was entirely eliminated in light-stimulated OptoSOS embryos, consistent with the reported repression of *sna* by *hkb* (Figure 1H)^12^. These data can be readily interpreted in the current model of terminal signaling, which holds that Erk signaling is necessary for PMG specification^13^. Our data suggests Erk is also sufficient to drive cells to adopt a contractile, PMG-like fate at all but the anterior-most positions in the embryo.

Erk’s ability to reshape cell fate responses raises important questions when considered in the context of normal embryogenesis. In wild-type embryos, Erk is normally activated to a high level in three distinct regions, yet only one (the PMG) contracts. At the anterior pole, the connection between Erk activation and tissue contractility appears to be strongly suppressed, as it is not observed in either wild-type or light-stimulated OptoSOS embryos. The situation is different along the ventral midline, where high levels of Erk activity are not normally associated with tissue invagination, but where optogenetic stimulation is sufficient to trigger contractility in these same cells. If Erk is sufficient to induce PMG specification, how is contractility normally prevented at the anterior pole and along the ventral midline?

We first turned our attention to the anterior pole, where even uniform OptoSOS stimulation fails to induce *fog/mist* expression or tissue contraction. We reasoned that contractility may be regulated by combinatorial control: high levels of the anterior-localized transcription factor Bicoid (Bcd) might block the Erk-dependent contraction program (Figure 2A). We generated embryos from OptoSOS mothers that are also homozygous for the *bcd^E1^* loss-of-function allele (termed OptoSOS-*bcd*) and imaged their gastrulation using differential interference contrast (DIC) microscopy. OptoSOS embryos exhibited tissue contraction across the posterior 85% of the embryo, leading to the flow of yolk toward the anterior pole and a thinning of the epithelial monolayer there (Figure 2B; Movie S4, top). In contrast, OptoSOS-*bcd* embryos exhibited uniform, isotropic contraction, suppressing virtually all overall tissue movements at gastrulation (Figure 2C; Movie S4, bottom). These embryos were still subjected to strong compressive forces, as many OptoSOS-*bcd* embryos popped, ejecting yolk and cells out of one or both poles (Movie S5). Thus, in the absence of Bicoid, Erk is sufficient to ‘posterior-ize’ the entire embryo, converting it into an apically-constricting epithelium but suppressing cell movements. The ability to induce invagination at any embryonic position in OptoSOS-*bcd* embryos will likely be useful in future studies for relating mechanical forces to tissue/embryo shape^14^.

**Figure 2.**
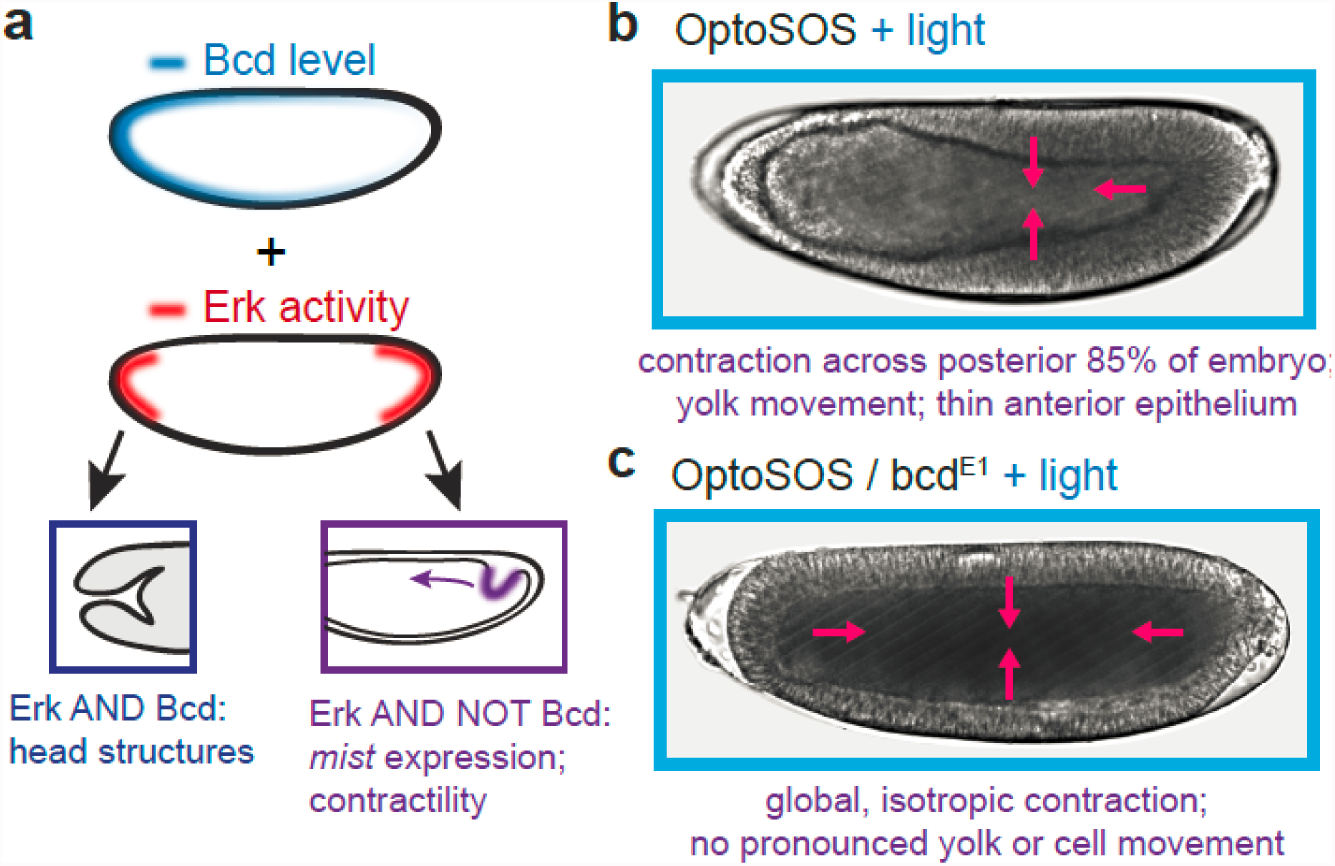
Erk-induced contractility is global and isotropic in the absence of Bicoid activity. (**a**) Schematic illustrating the hypothesis that anterior Bicoid activity is combinatorially interpreted with Erk to regulate tissue contractility. (**b-c**) DIC images of gastrulating (**b**) OptoSOS and (**c**) OptoSOS-*bcd* embryos. In **b** contraction does not extend to the anterior pole, where pressure induces epithelial thinning; in **c**, contraction is isotropic and blocks tissue reorganization.

We next turned to the ventral midline, where Erk normally programs non-contractile cell fates but where additional, ectopic Erk signaling is able to induce contraction. We hypothesized that Erk’s ability to trigger either of these responses from the same cell population is due to dynamic control. Such a model would hold that altering the timing or duration of Erk activity can select between distinct cell fates. Indeed, quantitative studies of Erk dynamics in wild-type embryos revealed that the timing and duration of Erk activity differs between the ventral midline and contractile posterior pole (Figure 3A)^15^. Posterior Erk signaling is initiated earlier, during the embryo’s first few nuclear cycles, whereas lateral Erk signaling is induced just prior to gastrulation during the 14^th^ nuclear cycle^15^. The duration of Erk signaling also differs: Erk is activated at the ventral midline in a transient 20 min pulse, whereas terminal signaling is sustained for over 1 h. Two models of dynamic control could be invoked to explain the lack of Erk-driven contractility in wild-type embryos. Contractility may be timing-dependent, such that cells are only able to produce *fog* and *mist* during the early time window when terminal signaling is active but ventral midline signaling is not. Alternatively, contractility may be duration-dependent, induced only in response to the long-term Erk signaling normally found at the poles.

**Figure 3.**
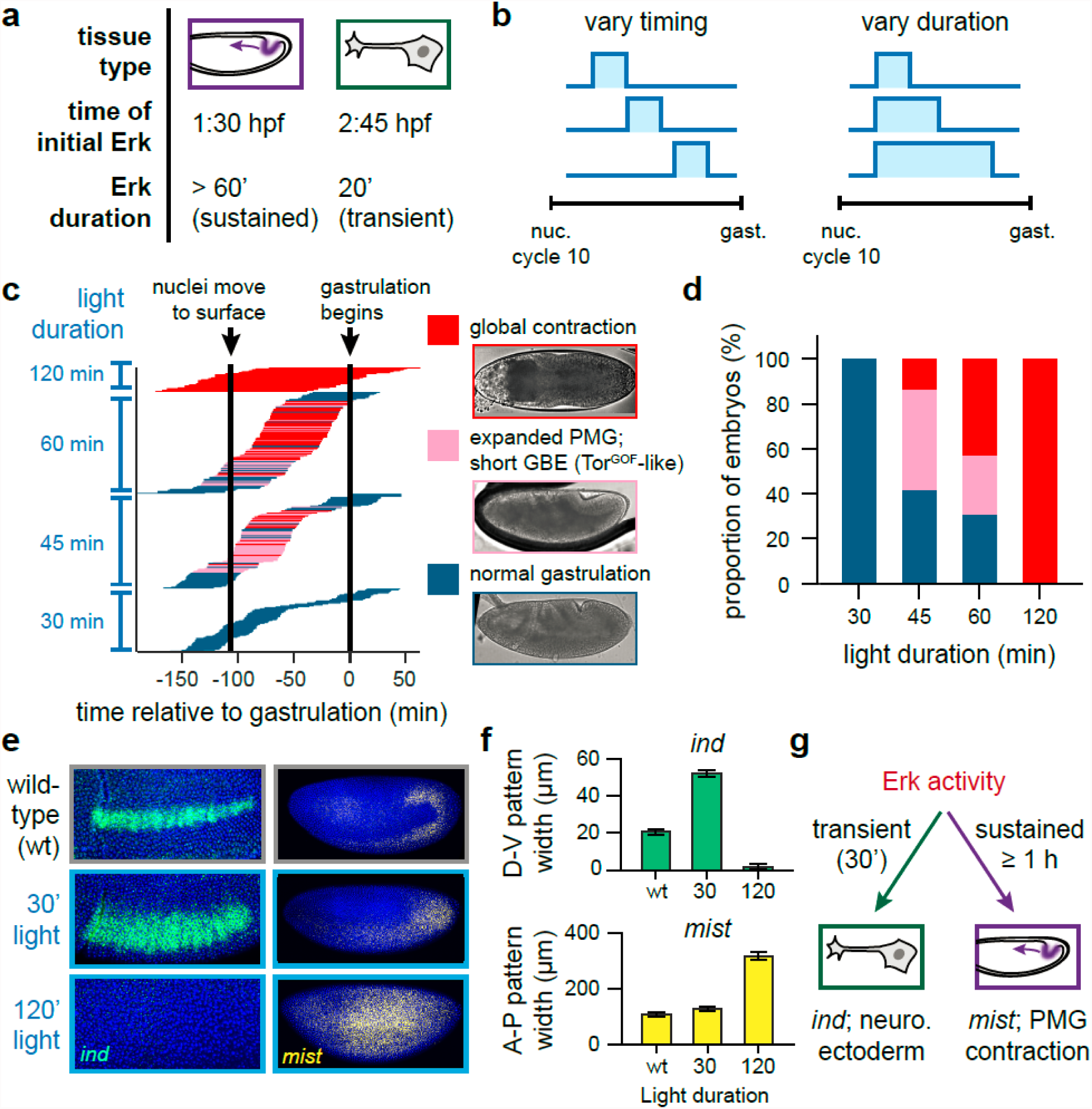
The duration of Erk activity controls a cell fate switch in the early Drosophila embryo. (**a**) Differences in the timing and duration of Erk activity may program posterior (left) and neurogenic fates (right). Posterior Erk signaling is sustained from NC10 to halfway through NC14, while Erk activity at the ventral midline is induced for approximately 20 minutes prior to gastrulation. (**b**) Optogenetic control can dissect whether timing or duration contributes to cell fate by varying each parameter independently. (**c**) Experimental data of embryo phenotypes for various times and durations of Erk activation. Each horizontal line represents one embryo. The length and position of the line represents the timing of light stimulation, while its color represents its gastrulation phenotype. (See also Figure S3). (**d**) Fraction of embryos exhibiting each gastrulation phenotype as a function of stimulus duration for all embryos in **c**. (**e**) Representative images of RNA FISH for *ind* (at the start of gastrulation) and *mist* (prior to gastrulation) for wild-type embryos (top) and OptoSOS embryos stimulated for 30 min or 2 h. (**e**) Quantification of the width of each expression pattern (D-V for *ind*; A-P for *mist*) for embryos stimulated as in **d**. (**f**) Model of how Erk-induced neurogenic and contractile fates are distinguished. Transient Erk activity gives rise to *ind* expression and neurogenic fates, while sustained activity programs *mist* expression and tissue contractility.

We set out to test both possibilities by systematically varying the timing and duration of light stimuli applied to individual OptoSOS embryos (Figure 3B-C). We stimulated 288 individual OptoSOS embryos of varying ages with either 30, 45, 60, or 120 min of blue light and imaged their progression through gastrulation using only far-red illumination to avoid introducing additional optogenetic stimulation (Figure S3A-B). Embryos subjected to a 30 min pulse of Erk activation exhibited normal gastrulation that was indistinguishable from control embryos (Figure 3C, bottom), although this brief stimulus triggered abdominal segment fusions similar to those induced by gain-of-function MEK mutants (Figure S3C). At the other extreme, stimulation with a 60 min or 120 min light pulse triggered a full contractile response (Figure 3C, top). Finally, a 45 min light pulse led to an intermediate phenotype, a partial expansion of the posterior domain similar to what is observed in Tor^GOF^ embryos (Figure 3C, middle; Figure S3D-E; Movie S6). This intermediate phenotype may be due to Erk signaling in the expanded domain being summed from our light input and the endogenous terminal pattern, which is initially broad and narrows to the poles over the first two hours of embryogenesis^16^.

These observations were highly informative about how Erk activity is interpreted into a contractile response. The same light intensity was used for all experiments yet led to outcomes varying from cuticle fusion to global contraction, allowing us to conclude that distinct Erk-dependent phenotypes do not arise simply from differences in the amplitude of Erk activation, as would be expected from a canonical morphogen. Varying the developmental time at which each light dose was delivered had no clear effect, ruling out the possibility that contractile cell fates are only established during a specific temporal window. In contrast, varying the duration of Erk activation was highly predictive of gastrulation and cuticle phenotypes (Figure 3D; Figure S4A). These data are suggestive of dynamic cell fate control, where differences in signal duration are sufficient to determine whether a ventral midRef16line or posterior cell fate is adopted.

If our dynamic control model is correct, short-duration Erk signaling should trigger the differentiation of additional neuronal precursors by increasing the size of the lateral domain where Erk and Dorsal are both present^15^. To test this prediction we stimulated OptoSOS cells with 0, 30 min or 120 min of light and stained for *intermediate neuroblasts defective* (*ind*), a gene that marks an Erk-dependent population of neuroblasts in the ventral midline, and *mist*, which marks contractile cells (Figure 3E-F). We found that stimulation with a transient, 30’ light pulse expanded the domain of *ind* expression dorsally without altering *mist* in those cells, consistent with the co-requirement for Erk and intermediate Dorsal levels to drive *ind* expression^17^. In contrast, 120’ of light abolished *ind* expression, instead inducing *mist* expression in lateral cells. Thus, varying a single dynamic feature – the duration of Erk stimulation – is sufficient to program cells on the lateral surface of the embryo to adopt distinct ventral midline or posterior cell fates (Figure 3G).

Differences in signaling dynamics have long been hypothesized to select among Erk-dependent cell fates^18, 19^, and at least two distinct mechanisms for decoding Erk dynamics have been proposed. The classic model is that of a persistence detector, where downstream genes sense the duration of a single Erk pulse through a combination of fast and slow regulatory steps (e.g. Erk-dependent transcription and post-translational regulation of gene products)^3, 20-22^. An alternative model is that of cumulative load detection, where distinct fates are triggered when the total integrated signal crosses a threshold^23^^-^^25^. These two types of dynamic decoding can be distinguished by their responses to continuous and discontinuous stimuli of the same duration (Figure 4A). A cumulative load model would predict identical responses regardless how the same total stimulus is divided into discrete pulses, whereas a persistence detector would only trigger a response to stimuli that are sustained continuously over time.

**Figure 4.**
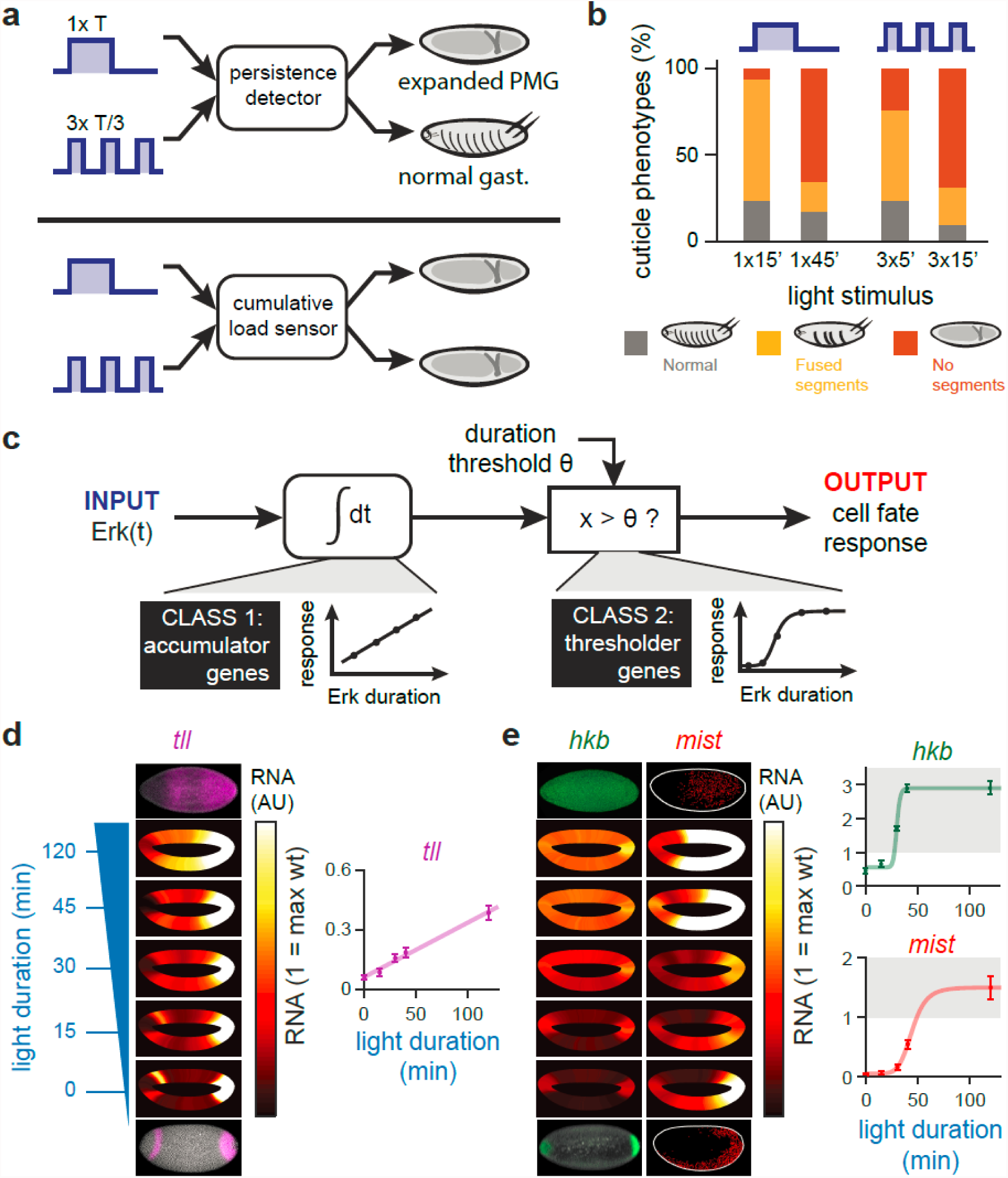
Embryo phenotypes are determined by the cumulative load of Erk signaling. (**a**) Two models of dynamic decoding – cumulative load sensing and persistence detection, can be discriminated by multiple-pulse stimulation. (**b**) Optogenetic embryos are exposed to a single 15 or 45 minute pulse of blue light (left) or an equivalent duration of light split into three equal pulses over a 90 minute window (right) and assayed for cuticle phenotypes. (**c**) Model of cumulative load sensing by an accumulator/ thresholder circuit. An accumulator node would increase linearly in response to Erk activity until a critical threshold *θ* is reached, at which point a thresholder node would turn on in a switch-like fashion. (**d,e**) Analysis of RNA FISH data from OptoSOS embryos exposed to varying durations of blue light and stained for (**d**) *tll* and (**e**) *hkb*/*mist* (see also Figure S4). The expression level for at least 50 embryos per condition is shown as an embryo-shaped heatmap (left) and the dorsal-most expression level is plotted as a function of duration of stimulation (right).

To determine whether Erk is sensed through cumulative load or persistence detection, we applied light as either a single bolus or as three pulses that were evenly spaced over a 90 minute window prior to gastrulation. We measured embryo responses by cuticle preparation, a sensitive and high-throughput assay for early embryonic phenotypes. Short-duration Erk activation leads to fused segments but do not disrupt gastrulation, whereas the major gastrulation defects produced by the contractile phenotype produce an empty cuticle (Figure S4A). Applying a single 15 minute Erk stimulus caused a majority of embryos to exhibit segment fusions, whereas a 45 minute Erk stimulus abolished segments entirely (Figure 4B; left bars). The same Erk doses divided into three equal pulses (3×5’ or 3×15’, respectively) led to similar phenotypes to those observed after single doses of the same total length (Figure 4B; right bars). Even though a single 45 min pulse was the minimum Erk dose that led to gastrulation failures (Figure 3C), three 15’ pulses were able to achieve the same phenotype, suggesting that the three-pulse regiment was interpreted by cells to be quite similar in magnitude. We also note that MEK and Tor gain-of-function mutants, which activate Erk to low, constant levels (Figure S1C), induce intermediate phenotypes that are quite similar to those driven by transient 30 min and 45 min light stimuli (Figure S3C-E). These results are consistent with an equivalence between long-term, low-amplitude signaling versus a brief, high-intensity pulse of Erk activity.

How might cumulative load trigger a cell fate switch? The simplest model requires two signal processing components, an ‘accumulator’ and ‘thresholder’ (Figure 4C). The accumulator component passively integrates Erk signal over time, providing memory of the total amount of Erk signaling that has been delivered. The thresholder would act downstream of the accumulator, comparing the accumulated signal to a fixed threshold and triggering a response only after the threshold is crossed. Such a model would predict two distinct behaviors among Erk-dependent genes: some may act as accumulators whose levels would rise in proportion to the total light input, whereas others would act as thresholders and would increase abruptly above a stimulus threshold.

If our cumulative load model were implemented by an Erk-responsive gene network, we would expect to find Erk target genes in both response classes. We thus stained for the terminal gap genes *tll* and *hkb* and the GPCR *mist* in response to different durations of light stimulation. We quantified RNA levels as a function of position in at least 50 embryos for each condition (Figure 4D,E; left). We then plotted RNA levels on the dorsal surface (a region where there was little accumulation in wild-type embryos) as a function of stimulus duration (Figure 4D,E; right). We found that *tll* responded as would be predicted for an accumulator gene: its levels linearly increased in proportion to the duration of Erk signal (Figure 4D). In contrast, *hkb* and *mist* acted as threshold genes, turning on abruptly at 30 min and 45 min of signaling, respectively (Figure 4E). Notably, the threshold duration at which *mist* is switched on coincides exactly with the threshold for large-scale tissue contractility, consistent with *mist*’s role in this process. Our results demonstrate that distinct Erk target genes exhibit qualitatively different response functions, some acting as linear signal integrators and others as all-or-none switches above a threshold stimulus level. These observations are consistent with our model that the cumulative load of Erk signaling, not the duration of a single persistent pulse, is decoded into distinct cell fates in the early *Drosophila* embryo.

Through a combination of genetic perturbations and time-varying optogenetic stimuli, the current study begins to define a model for how Erk is capable of programming at least three distinct cell fates the early embryo (Figure 5). At the posterior, sustained Erk signaling induces expression of the Fog/Mist receptor-ligand pair, leading to apical constriction and tissue invagination. At the anterior, contractility is suppressed by Bcd, whose combination with Erk leads to anterior-specific gene expression. Finally, in the middle of the embryo, transient Erk activation induces the formation of neuroblasts, a fate that can be overridden by additional, ectopic Erk activity.

**Figure 5.**
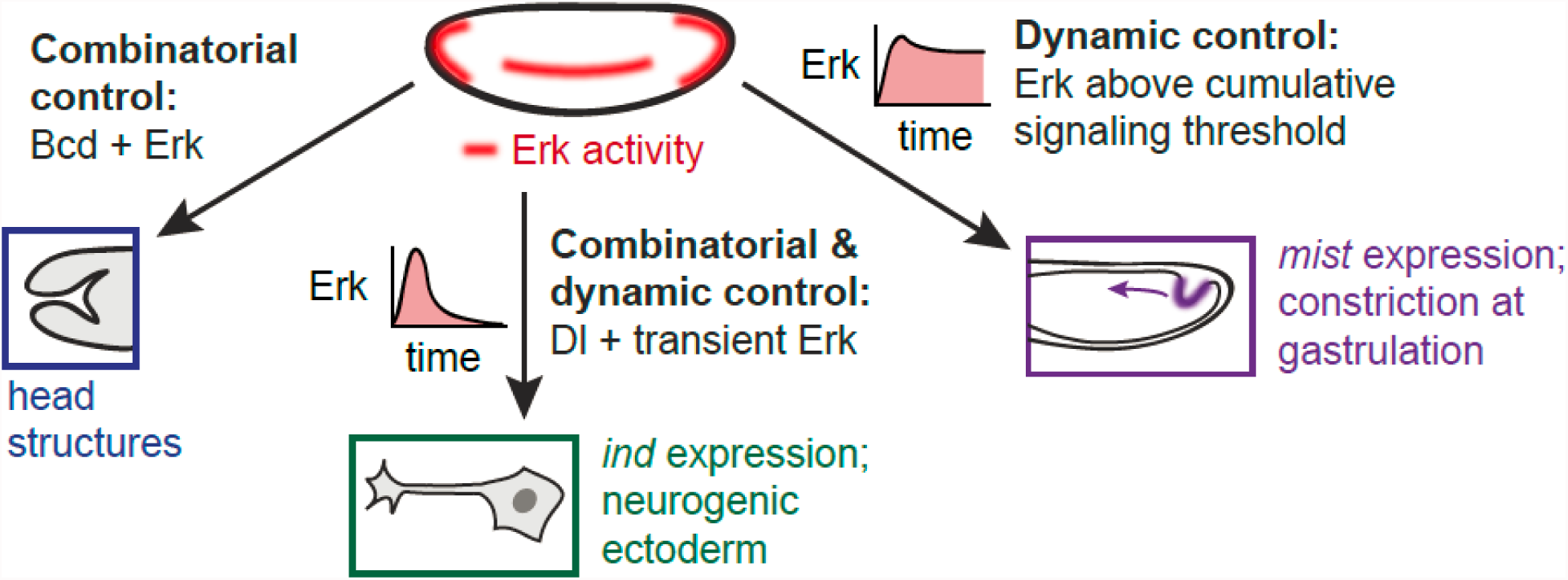
Conceptual model of Erk-dependent cell fate control in the early embryo. At the anterior pole, combinatorial control with Bicoid prevent contraction and specify head structures. Along the ventral midline, transient Erk activity elicits neurogenic fates, whereas sustained Erk activity in the posterior induces posterior midgut differentiation, contraction at gastrulation, and suppression of neurogenic fates. In this model, dynamic control is sufficient to distinguish between PMG and neurogenic fates in embryos exposed to intermediate levels of Dorsal.

In contrast to the now-classic model for how Erk dynamics are decoded in cultured cells^3, 21, 26^ we find that the early *Drosophila* embryo senses the cumulative load of Erk signaling rather than the length of a persistent stimulus. This distinction is important, as the putative network architectures that perform these two signal processing functions can be quite different. Persistence detection is thought to rely on network motifs like the coherent feedforward loop^27^, whereas cumulative load detection can be implemented by combining long-term integration with an ultrasensitive downstream step. Elucidating the dynamic input-output response of a biological network can be a crucial first step towards a complete understanding of its network architecture and molecular components.

To our knowledge, this study is the first direct demonstration that Erk dynamics control cell fate *in vivo*, despite the phenomenon’s long history of study in cultured cells. We would argue that the early *Drosophila* embryo has all of the characteristics of an ideal model system: cell fates are specified within 3 hours and are highly reproducible between embryos, Erk-dependent tissue contractility can be observed directly by brightfield microscopy, the molecular candidates for decoding dynamics are limited to a few active zygotic genes in the early embryo, and the combination of optogenetic and classical genetic tools enable complex perturbations of these network components. It is also likely to be only one of many *in vivo* examples of dynamic control, suggesting that optogenetics could play a major role in future studies for uncovering and dissecting complex dynamics-dependent cell fate choices.

## Author Contributions

H.E.J., S.Y.S. and J.E.T. conceived and designed the project. H.E.J. performed all experiments. H.E.J. and J.E.T. wrote the manuscript with input from all authors.

## Acknowledgements

We thank all members of the Toettcher lab for helpful comments, as well as Eric Wieschaus for his suggestions and insights throughout the study. Yuji Yamazaki kindly provided the *mist* RNA probe. HEJ was supported by NIH Ruth Kirschstein fellowship F32GM119297. This work was also supported by NIH grant DP2EB024247 and NSF CAREER Award 1750663 (to J.E.T.), and NIH grant R01GM086537 (SYS). We also thank Dr. Gary Laevsky and the Molecular Biology Microscopy Core, which is a Nikon Center of Excellence, for microscopy support.

